# Oculomotor nerve requires an early interaction with muscle precursors for nerve guidance and branch patterning

**DOI:** 10.1101/483396

**Authors:** Brielle Bjorke, Katherine G. Weller, G. Eric Robinson, Michelle Vesser, Lisheng Chen, Philip J. Gage, Thomas W. Gould, Grant S. Mastick

**Affiliations:** Department of Biology, University of Nevada, Reno, NV 89557 USA; Department of Ophthalmology & Visual Science, University of Michigan Medical School, Ann Arbor, MI 48105, USA; Department of Physiology and Cell Biology, University of Nevada School of Medicine, Reno, United States

**Keywords:** Oculomotor nerve, axon guidance, extraocular muscle, muscle precursor, neuromuscular targeting

## Abstract

Muscle function is dependent on innervation by the correct motor nerves. Motor nerves are composed of motor axons that extend through peripheral tissues as a compact bundle, but then diverge to create nerve branches to specific muscle targets. A transition point typically occurs as motor nerves grow near their targets, where the fasciculated nerve halts further growth, then later initiates branching to muscles. The motor nerve transition point is potentially an intermediate target acting as a guidepost to present specific cellular and molecular signals for navigation. Here we describe the navigation of the oculomotor nerve with respect to eye muscle precursor cells in mouse embryos. We found that the oculomotor nerve initially grew to the eye three days prior to the appearance of any eye muscles. The oculomotor axons spread to form a plexus within a mass of eye muscle precursors, then the nerve growth paused for more than two days. This plexus persisted during primary extraocular myogenesis, with a subsequent phase in which the nerve branched out to specific muscles. To test the functional significance of the nerve-precursor contact in the plexus, we genetically ablated muscle precursors early in nerve development, prior to nerve contact. Ablation of muscle precursors resulted in oculomotor nerve fibers failing to stop to form the plexus, but instead growing past the eye. In contrast, ablating the precursor pool at later stages, after the nerve has contacted the precursor cells, results in ectopic branching restricted near the eye. These results demonstrate that muscle precursors act as an intermediate target for nerve guidance, and are required for the oculomotor nerve to transition between nerve growth and distinct stages of terminal axon branching.

## Introduction

During embryonic development, motor axons project out from the spinal cord and brain stem into peripheral tissues as large fasciculated nerves that supply a limb or organ. Upon reaching a peripheral target region, specific sub-populations of motor axons must halt further growth, defasciculate from the main nerve trunk and reorient as nerve branches toward specific muscles. Although guidance cues have been identified that act in initial motor axon guidance, little is known about the cues and tissues that guide nerve transition between growth and branching to specific muscles.

An example of such a transition occurs during the course of oculomotor nerve development. The oculomotor nerve, along with the trochlear and abducens nerve, innervate the extraocular muscles. During development in mouse and chick embryos, oculomotor axons extend from the ventral midbrain toward the developing eye, and upon reaching a ‘peripheral decision region’, reorient to form specific branches toward four extraocular muscle targets (Cheng et al., 2014; Michalak et al., 2017). Similarly in zebrafish, oculomotor axon growth cones transition from exploratory behavior to axon subgroups that align to project to individual muscles (Clark et al., 2013). Interestingly, several human mutations that result in eye muscle mis-innervation cause developmental oculomotor nerve errors in the ‘peripheral decision region’ in animal models (Chilton and Guthrie, 2017; Clark et al., 2013; Park et al., 2016; Whitman and Engle, 2017)

The tissue substrates for the extending oculomotor nerve are likely important for guidance and branching decisions. However, to date, no substrate capable of influencing nerve growth and branching prior to muscle formation has been described for the oculomotor nerve. Historic studies in chick describe a mass of cells located at the tip of the developing nerve (Carpenter, 1906), potentially coinciding with the ‘peripheral decision region’, although the identity of this cell mass has not been molecularly defined. Similarly, spinal nerves pause when they reach a mass of mesodermal cells prior to entering the limb (Hollyday and Morgan-Carr, 1995; Tosney and Landmesser, 1985a, b; Wang and Scott, 2000). The mass of spinal mesodermal cells coincides with a decision region where axons pause before making distinct pathway choices (Lance-Jones and Landmesser, 1981; Tosney and Landmesser, 1985a, b). In grasshopper and fly, motor nerves interact with muscle precursor cells (Ball et al., 1985; Ho et al., 1983). Functional tests in grasshopper show that, a single muscle precursor is sufficient to cause an entire nerve to branch (Landgraf et al., 1999), and the removal of muscle precursors results in motor nerves that do not branch (Prokop et al., 1996). Similarly, zebrafish, muscle precursors act as intermediate targets that halt motor axon extension (Melancon et al., 1997). Together these findings suggest that growing motor nerves may be guided by contact with muscle precursors.

A recent study, published while our manuscript was being prepared, described mouse oculomotor nerve development patterns through the stages of nerve innervation of the eye muscles (Michalak et al., 2017). This study also ablated eye muscles using Myf^Cre^ mutant embryos, finding that the oculomotor nerve projects normally out to the eye, but then the major superior branch fails to form, inferior sub-branches fail to form terminal arbors in the absence of their targets, and that the oculomotor neurons died within several days (Michalak et al., 2017). However, this study did not look early enough to see the initial outgrowth of pioneer axons of the oculomotor nerve, or examine potential contacts with eye muscle precursor cells. In the developing oculomotor system, the transcription factor Pitx2 is an early marker for muscle precursor cells that differentiate to form the extraocular muscles (EOMs) (Zacharias et al., 2011). Pitx2+ precursor cells form a thick wedge in the mesenchyme ventrolateral to the eye early in development (Zacharias et al., 2011). This suggests that a mass of eye muscle progenitor cells may define the ‘peripheral decision region’ for the oculomotor nerve. In our present study, we sought to determine whether Pitx2-positive muscle precursors regulate oculomotor nerve pathfinding, branching and innervation by acting as a guidepost at the peripheral decision region. To test this hypothesis, we examined the development of this nerve in response to early or late muscle precursor ablation. Our data demonstrate that oculomotor nerve growth and branching require interaction with muscle precursor cells.

## Materials and Methods

### Mouse embryos

Animal experiments were approved by the UNR IACUC, following NIH guidelines. Wild type CD-1 mice were purchased from Charles River Laboratories (Wilmington, MA USA). Pitx2 mutant and control litter mate embryos were produced by crossing Pitx2 heterozygotes (Zacharias et al., 2011). Myf5-DTA embryos were obtained by crossing a Myf5-Cre line (JAX B6.129S4-Myf5^tm3(cre)Sor/J^) (Tallquist et al., 2000) to a second transgenic floxed DTA line (JAX stock #006331, Gt(ROSA)26Sor^tm1(DTA)Jpmb^) (Ivanova et al., 2005). Embryos were genotyped by PCR against the Cre and DTA genes. Embryos were collected and fixed in 4% PFA in 0.1M PO4 buffer, as described in (Bjorke et al., 2016). Embryos were then either labeled as whole mounts by antibody labeling or axon tracing, or were embedded for cryosectioning and antibody labeling. For each label type and embryonic stage, the data shown is representative of three (or more) biological replicates.

### Immunohistochemistry

Antibody labeling was as described in (Farmer et al., 2008). Primary antibodies included Pitx2 (1:1000, rabbit anti-rodent/human Pitx2 peptide antigen, Capra Science), MyoD (Dako, clone 5.8A), MyoHC (1:5, MF 20, Developmental Studies Hybridoma Bank), beta-III-tubulin (1:2000, TuJ1, mouse anti-rat beta-III-tubulin, Biolegend). The Pitx2 antibody labeling pattern was validated by noting matching expression in eye muscle precursors as seen by an independent Pitx2 antibody reported by Zacharias, et al. (2011), and the specificity of that antibody was confirmed by noting a complete loss of labeling in sections of Pitx2 null embryos (PJG, data not shown). Secondary antibodies included anti-rabbit and anti-mouse Alexa 488 and Alexa 555 (Invitrogen).

### Axon tracing

To label the oculomotor nerve, the lipophilic dye, DiI (Invitrogen/Thermo Fisher Scientific) was crushed onto the oculomotor nerve, as described (Bjorke et al., 2016). The dye tracer was allowed to diffuse by incubation in 4% PFA for 1 day at 37 degrees Labeled embryos were then cleared in 50% glycerol for a few hours before mounting under a coverslip for imaging. Images were obtained by combining confocal images via Z-stack on a Leica SP8 confocal microscope. Data shown are representative images of the stated numbers of embryos labeled; Investigators were not blinded to the embryo genotypes; Labels were considered successful and included in the analysis if the diI tracer reached the end of the axons (growth cones), and excluded if not. Unfortunately, the incubation at high temperature to allow diI transport prevented subsequent antibody labeling, such as to label other nerve or muscle markers.

## Results and Discussion

### The oculomotor nerve initially grows to a mass of muscle precursor cells next to the eye

To investigate whether muscle precursors are in position to play a role in the transition between nerve growth and branching, we labeled motor nerves and muscle precursors during initial nerve development from E9.5-14.5. Motor nerves were labeled by driving GFP under the Islet promoter (Lewcock et al., 2007), and muscle precursors were labeled immunolabeled against the transcription factor Pitx2. On E9.5, motor axons began to exit from the oculomotor nucleus in the ventral midbrain into peripheral mesenchyme within the cephalic flexure. The initial nerve navigated parallel to the ventral midline of the forebrain. On E9.5, no Pitx2 protein was found in the mesenchyme (**Fig. 1A**). However, by E10.0, oculomotor axons reached an area ventrolateral to the eye that was diffusely populated with Pitx2+ cells (**Fig. 1A**). Thus, the first target of the oculomotor nerve is Pitx2+ precursor cells, not the eye itself or mature muscle fibers. These Pitx2+ cells condensed to form a wedge shape on E11.0 but maintained contact with the now thick fasciculated oculomotor nerve. The nerve fanned out at the tip to give the appearance of a plexus within the mass of Pitx2+ precursor cells (**Fig. 1C,D**). Scanning the length and depth of the Pitx2+ mass, we found that two smaller nerve plexuses neighbored the oculomotor nerve plexus, although arriving about a day after the oculomotor nerve. By following the trajectories of these nerves between sections of E11.5 and 12.5 embryos, we identified them as the trochlear nerve (originating from r1 isthmus) and abducens (originating from r5/6). The fasciculated abducens nerve terminated ventrolateral and deep to the oculomotor plexus, while the less fasciculated trochlear nerve contacted precursors dorsal to the oculomotor plexus (**Fig. 1E,F**).

**Figure 1.**
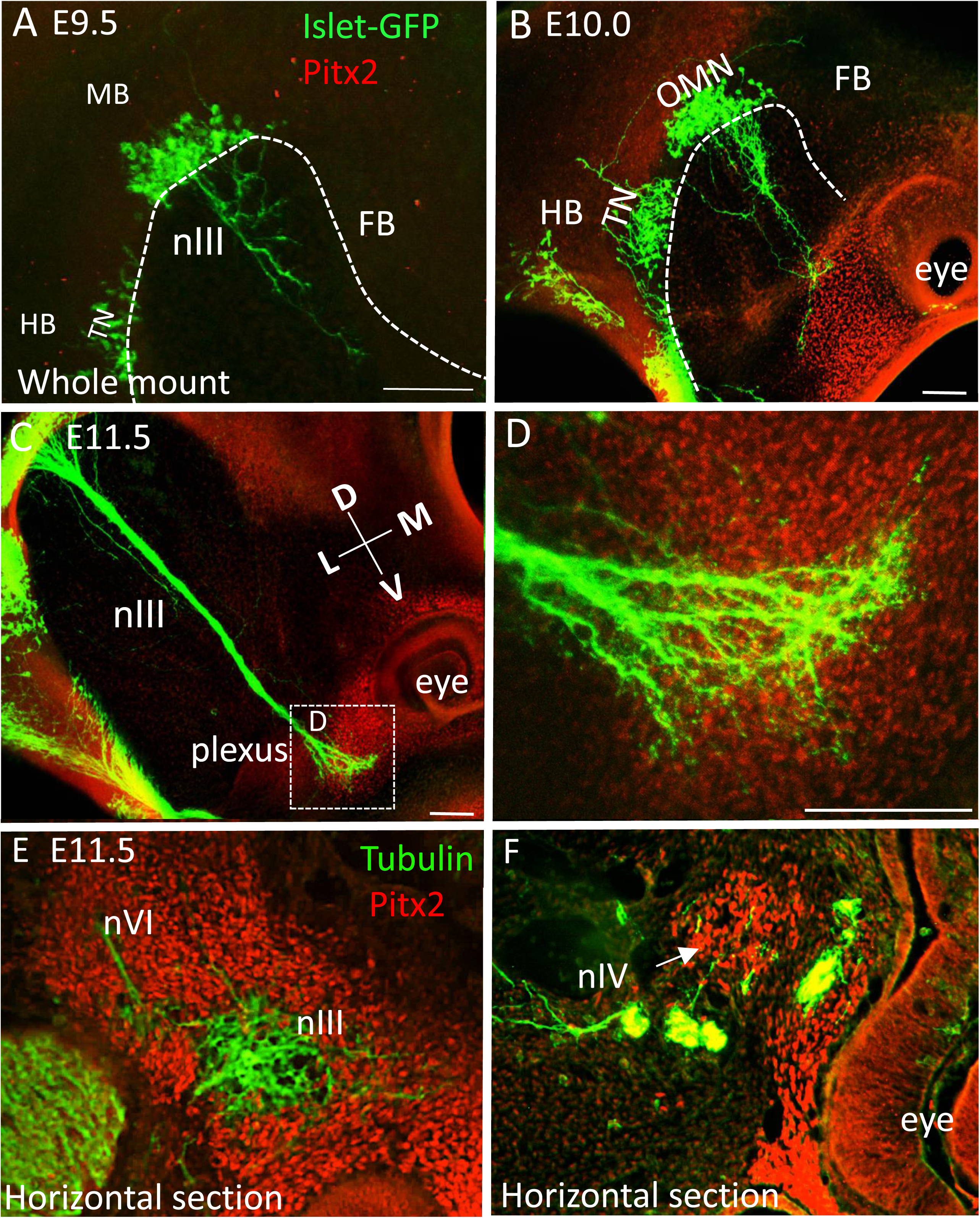
The initial developmental target of the oculomotor nerve is a mass of Pitx2+ precursor cells. The developing oculomotor nerve projections were compared with the location of muscle precursor cells in both whole mounts (**A-D**) and sections (**E,F**). Motor neurons were labeled with GFP driven by the Islet promoter (green), and muscle and tendinous precursor cells labeled with an antibody against the transcription factor Pitx2 (red). **A.** On E9.5, oculomotor nerve fibers projected out of the ventral neural tube into peripheral mesenchymal tissue toward the eye. **B.** On E10.0, oculomotor nerve fibers made first contact with Pitx2+ cells ventrolateral to the eye. **C,** On E11.5, oculomotor nerve fibers fasciculated into a compact nerve bundle. At the tip, the nerve fanned out to form a plexus embedded within the mass of Pitx2+ precursor cells. **E.** The trochlear nerve formed a plexus in a distinct area ventrolateral and deep to the oculomotor plexus. **F.** The abducens nerve fibers interacted with Pitx2+ precursors dorsal to the oculomotor plexus. Abbreviations: nIII, oculomotor nerve; TN, trochlear nIV nucleus; MB, midbrain; HB, hindbrain; FB, forebrain; D, dorsal; V, ventral; L, lateral; M, medial; nVI, abduscens nerve; nIV, trochlear nerve. Each image shown is representative of successful labels of 3 (or more) biological replicates. Scale bar, 100 µm.

Therefore, muscle precursors cells are an intermediate target of the three nerves that innervate the extraocular muscles. Differing points of oculomotor, trochlear and abducens nerve contact with the precursor mass suggest that the nerves can recognize sub-regions within the precursor mass. The precocious contact of motor nerves with muscle precursors is consistent with a prior study of the abducens nerve, in which the nerve branched to contact distinct subsets of mesenchymal condensations, presumed for lateral rectus and other facial muscles (Wahl et al., 1994). Our labels extend this by showing that the oculomotor nerves extend to contact Pitx2-expressing muscle precursor masses. This pre-patterning of the muscle precursor mass potentially guides the matching of nerves to muscle precursors via distinct secreted or contact-mediated cues, which remain unknown.

### Axons of the developing oculomotor nerve stall in the muscle precursor plexus during the period of primary myogenesis

Extraocular muscles are derived from prechordal mesoderm and cranial paraxial mesoderm (Gage et al., 2005; Lescroart et al., 2010; Noden and Francis-West, 2006), and like typical somite-derived trunk muscles, extraocular muscles undergo a developmental pattern of primary myogenesis, in which individual muscle precursors differentiate to myocytes that fuse to form multinucleated myotubes. A close association between the oculomotor nerve and muscle precursors prior to primary myogenesis suggests that the nerve may maintain contact with precursors as they progress through development, such that distinct subsets of axons may maintain an association with distinct precursors as they differentiate into distinct muscles. Surprisingly, we found that the oculomotor nerve did not maintain contact with precursor cells streaming around the eye or undergoing myogenesis defined by the appearance of myofibers (**Fig.2)**. Instead, the oculomotor nerve maintained a plexus in the central location with Pitx2+ cells, but that was devoid of myofibers as marked by Myosin Heavy Chain. (**Fig. 2A,B**). The nerve did form small dorsal axon projections on E12.5 to initiate the superior branch of the oculomotor nerve (arrow in **Fig.2B**). Likewise, a thicker inferior branch emerged (stubby arrow in **Fig. 2B**). Despite the initial formation of the superior and inferior branches, these nerves did not yet contact the nascent myofibers on E12.5 (**Fig. 2** C, n=4). By E13.0, the EOM myofibers established their stereotypical coordinate positions around the eye (**Fig.2 D,E**), extending from the eye to the orbit. However, only the superior rectus was contacted by axons (**Fig. 2D-F**, n=5). Nerve invasion into the remaining muscles was found on E13.5 (Fig. G-I, n=5), and occurred at the belly of each muscle. This timing was verified by applying a lipophilic dye to the oculomotor nerve on E12.5 and E13.5 (**Fig. 2J,K**). We observed that an adult innervation pattern is achieved on E14.5 in whole mount embryos viewed from the behind the eye (**Fig. 2L**).

**Figure 2.**
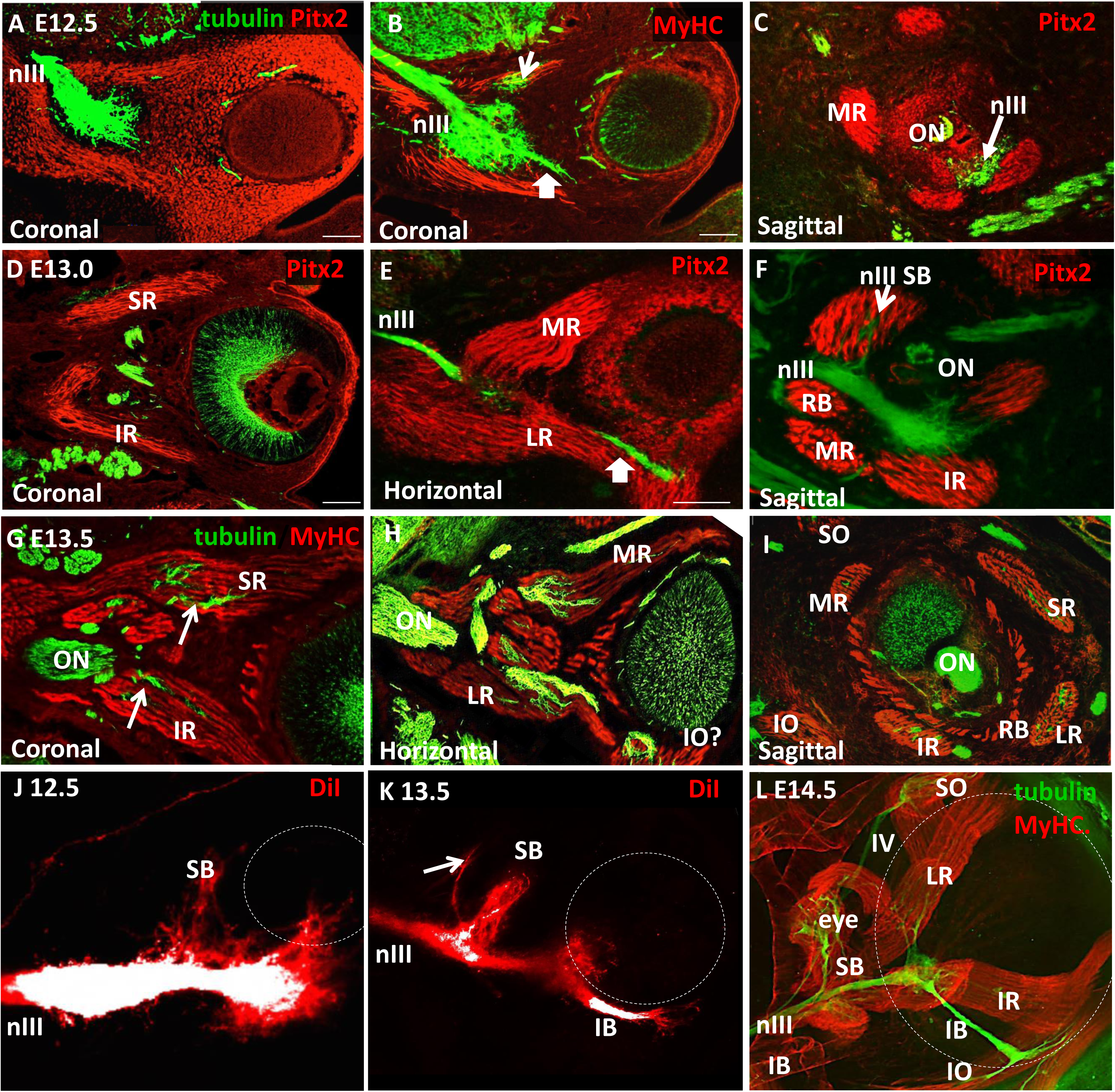
Oculomotor nerve fibers, centrally located in the Pitx2+ precursor mass, wait for extraocular muscle myogenesis before branching to innervate distinct muscle targets. (**A-I**) To characterize the timing and interaction between oculomotor nerve fibers and developing muscle fibers, we immunolabeled sections through the developing eye with βIII–tubulin (nerve, green), and either Pitx2 or Myosin Heavy Chain (MyHC, muscle cells, red). Myogenesis was determined by the appearance of myofibers that continue to express Pitx2, or by activation of MyHC in fibers. **A-C.** In coronal sections on E12.5, the nerve maintained a plexus centrally located within the Pitx2+ precursor mass, while precursor cells along the periphery of the mass began to stream to positions around the eye, followed by fiber formation and Myosin heavy chain expression. **B.** On E12.5, a small inferior branch of the oculomotor nerve could be seen projecting ventrally (arrow). **C.** A representative E12.5 sagittal section through extraocular muscles with no nerve fibers found within the developing muscles. ON, optic nerve; MR, medial rectus. **D-F.** On E13.0, most Pitx2 precursor cells fused to form distinct muscle fibers located around the eye. However, all muscles were devoid of nerve fibers except the superior rectus (SR) which was innervated by the superior branch (SB) of the oculomotor nerve. IR, inferior rectus; LR, lateral rectus; RB, retrobulbar. **G-I.** By E13.5, nerve fibers grew to contact each of the muscles. IO, inferior oblique; SO, superior oblique. **J, K.** Whole mount DiI tracing of the oculomotor nerve revealed a similar timeline of initial nerve branching toward the superior rectus muscle on E12.5, and a more developed superior (SB) and inferior (IB) division of the nerve on E13.5. The DiI label images were over-exposed in the main nerve branches (white) to reveal the smaller branches (red). **L.** In E14.5 whole mount immunolabeled preparations viewed from behind the eye, a mature innervation pattern is apparent, with each muscle receiving innervation. Each image shown is representative of successful labels of 3 (or more) biological replicates. Scale bars, 100 µm.

Taken together, these observations show that the oculomotor nerve pauses to make prolonged contact with muscle precursor cells, and that the nerve plexus is maintained during primary myogenesis. **Figure 3** shows a summary of the time course of the development of the oculomotor nerves, including nerve growth, positions of muscle precursors, and differentiation of muscles.

**Figure 3.**
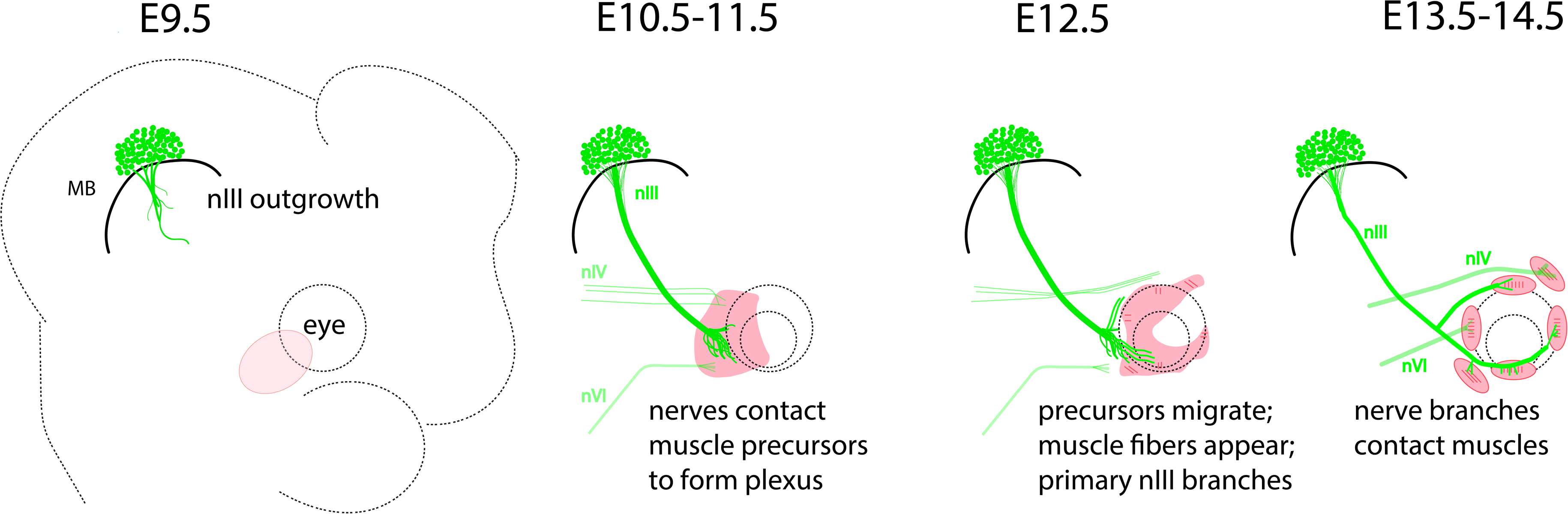
Time course of oculomotor development in mouse embryos. Schematic summarizing the main events in the development of the oculomotor nerves and extraocular muscles. The oculomotor nerve, nIII, is the first motor nerve to exit the CNS, from the midbrain (MB) on E9.5. Before E10.5, the nIII nerve contacts a mass of Pitx2 muscle precursors (pink) at the ventro-lateral side of the eye (dashed circles), with the axons spreading to form a plexus. The trochlear nIV and abduscens nVI nerves arrive a day later to contact separate areas within the muscle precursor mass. On E12.5, the muscle precursors migrate around the eye and begin to form muscle fibers at coordinate positions around the eye, while the main bifurcation of the oculomotor nerve begins to form the superior and inferior branches. On E13.5, the main branches of the oculomotor nerve contact and penetrate the muscles, completing this process by E14.5. (The cartoon disregards the rapid growth in embryo and eye size during this time course.)

### Muscle precursors provide a stop signal to the oculomotor nerve and are required for accurate nerve branching

The time course of oculomotor development demonstrates that the nerve interacts with precursor cells at a critical juncture: when the nerve fibers pause further growth, awaits primary myogenesis, and then resumes growth to branch and innervate specific muscles. The early and prolonged contact between the nerve plexus and the muscle precursor cells suggests that the precursor mass is positioned to regulate nerve growth and time nerve branching. To determine the functional significance of the early nerve interaction with the precursor mass, we used two genetic approaches: the first genetically ablated the precursors in Pitx2 mutant mice; the second reduced the number of precursor cells by ablation via driving diphtheria toxin under the Myf5 promoter (Myf5-DTA).

To completely remove muscle precursors and eliminate the cells that develop into extraocular muscles, we examined the oculomotor nerve projection in Pitx2 mutant embryos. Pitx2 mutant embryos lack EOMs due to precursor cell death just prior to E10.5 (Zacharias et al., 2011). We first confirmed that extraocular muscles did not develop in Pitx2 mutant embryos by labeling E11.0 embryos with Pitx2 and 13.5 embryos with MyHC. Both stages lacked Pitx2+, and MyHC + cells ventrolateral to the eye (data not shown), confirming the prior study (Zacharias et al., 2011).

To examine oculomotor nerve projection patterns, because the mutant mouse lines did not have the genetic motor axon transgene, we used diI tracing of the oculomotor nerve. This tracing strategy had the advantage of labeling a specific nerve, but the required high temperature incubation unfortunately prevented muscle antibody labeling. Control Pitx2+/+ embryos showed stereotypical superior and inferior branches of the oculomotor nerve ventrolateral to the eye (**SB, IB in Fig. 4A**). However, in Pitx2 homozygous mutant embryos, the oculomotor nerve reached the ventral aspect of the eye, but failed to pause and instead extended long branches past the eye **(Fig. 4B-D,** n=7). Interestingly, nerve projection patterns were not bilaterally symmetric, with different errors occurring on the left and right sides, suggesting that axon growth and guidance in the absence of muscle precursors is randomly determined around each eye. This further implies that once axons have been attracted to the vicinity of the eye, any remnant or redundant cues from the eye or nearby tissues are relatively weak. The strong overgrowth errors suggests that the precursor mass provides the signals to pause inappropriate further growth.

**Figure 4.**
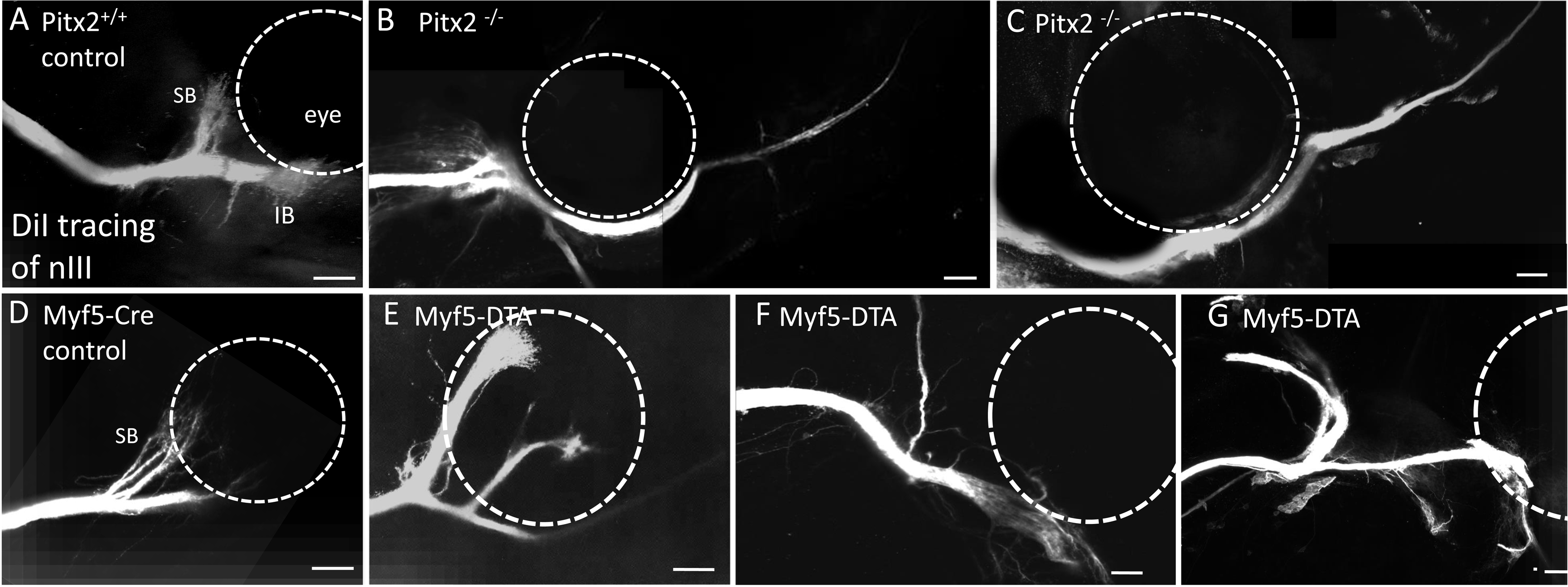
Extraocular muscle precursors are required to halt nerve growth and guide nerve branching. To examine the function of nerve contact with eye muscle precursors, we traced oculomotor nerve projections with diI on E12.5, in mouse embryos either lacking in extraocular muscle precursors in Pitx2 mutants (**A-D**), or having a severe reduction in extraocular precursors as described for Myf5-DTA mice (**D-H**). **A.** In Pitx2 homozygous wild type embryos, muscle precursor cells were left intact and a distinct inferior (IB) and superior branch (SB) was apparent ventrolateral to the eye. **B, C.** In Pitx2 homozygous mutant embryos that lacked muscle precursors, the nerve navigated to the ventrolateral region and along the ventral side of the eye, but then shot far past the eye in most cases (n= 5 out of 7 embryos, representing independent biological replicates). Side branches were absent (n=6 out of 7)(C), and the main nerve bundle projected abnormally past the eye (B, C). **D-G.** In Myf5-DTA mouse embryos that expressed diphtheria toxin under the Myf5 promoter, the main trunk of the oculomotor nerve generally did not overshoot the eye (n=5), but displayed large random branch patterns generally dorsal to the main nerve, including abnormal branches before the eye, and around the eye. In one case, a small branch extended past the eye (E). Control embryos (n=3) showed the expected initial bifurcation into relatively short and superior and inferior branches. Scale bar, 100 µm.

The severe overshooting of the oculomotor nerve contrasts with a similar recent study of embryos mutant for Myf5, a transcription factor required for extraocular muscle differentiation to ablate EOM (Michalak et al., 2017). Similar to our findings, the oculomotor nerve grew to the eye in the normal trajectory, then terminated in relatively short abnormal branches confined to the vicinity of the eye.

The more severe defects in Pitx2 mutants could be explained by a relativelylate timing of EOM primordia cell death on E11.5 in *Myf5* mutants, instead of the earlier E10.5 in Pitx2 mutants (Sambasivan et al., 2009; Zacharias et al., 2011). This suggests that the EOM precursors provide an early stop signal on E10.5 to inhibit further growth of the oculomotor nerve. Lack of a stop signal would result in the nerve overshooting phenotype found in Pitx2 mutants.

Our wild type time course demonstrates that E12.5 is a critical time point in oculomotor development when the nerve initiates branching toward the final eye muscle locations. To investigate whether the precursor mass influences nerve branching, we took advantage of the delayed expression of Myf5, relative to Pitx2, by expressing diphtheria toxin under the Myf5 promoter in Myf5^CreSOR^, R26^DTA^ mice (Haldar et al., 2007; Haldar et al., 2008). However, despite expression of DTA in all EOM precursors, some ‘escaper cells’ evade death and subsequently proliferate to eventually generate delayed extraocular muscles (Comai et al., 2014; Haldar et al., 2008). Therefore, driving diphtheria toxin under the myf5 promoter provides a unique opportunity to investigate how the nerve behaves when the precursor numbers are reduced on E11-E12.

Like Pitx2 mutant embryos, diI tracing of the oculomotor nerve in Myf5-DTA mutants showed nerves that reach the ventrolateral aspect of the eye. However, unlike the Pitx2 mutants, in Myf5-DTA embryos the main trunk of the oculomotor nerve did not shoot past the eye **(Figure 4D-G,** n=5). Instead, the nerve stopped near the ventrolateral aspect of the eye and either branched erratically around the eye with some branches shooting past the eye (**Figure 4E, n=1)** or formed a stubby terminal at the base of the eye, with one main branch that traversed dorsally in the area typical of a normally forming superior branch (**Figure 4F,G,** n=3 out of 5). These results suggest that muscle precursors have an additional function to organize terminal nerve branching.

We speculate that branch patterns may require changes in the expression of axon-axon adhesion molecules to axon-muscle adhesion molecules, and may also generate a change within the axons to express receptors to respond to muscle and mesenchyme derived guidance cues such as CXCL12, HGF and Sema3A and 3C (Chilton and Guthrie, 2004; Ferrario et al., 2012; Lerner et al., 2010). Our data suggests that these changes may be driven by signals from newly fused myocytes or myofibers. Future research may investigate the expression of adhesion molecules and guidance receptors expressed by motor axons in the plexus, and the timing of guidance cue expression in developing extraocular muscles.

Both our study, and Michalak et al. find that the oculomotor nerve reaches the ventrolateral aspect of the eye in embryos that fail to form extraocular muscles. The fidelity of the oculomotor projection suggests that extraocular muscle precursors are not required to attract the oculomotor nerve to the eye, because the nerve fibers navigate correctly when the muscles are ablated. However, the oculomotor nerve reaches the muscle precursors on E10.0, therefore eliminating these cells on E10.5 does not rule out the possibility that the muscle precursors provide attractive signals during the earlier period of E9-E10. We were unable to detect Pitx2 antibody labeling prior to E10.0 in the cranial mesenchyme. However Pitx2-driven LacZ expression is broadly expressed across the cranial mesoderm as early as E8.5 (Shih et al., 2007). Therefore, cells fated to become EOMs may play some early role in initial nerve guidance. Previous research has identified the expression of several cues expressed in and around the developing chick eye (Ferrario et al., 2012; Lerner et al., 2010; Ojeda et al., 2013). Future research may investigate whether these cues mediate muscle precursor influences on motor axons.

## Clinical significance of early muscle precursor interaction

In the last decade, there have been significant advances in elucidating the mechanisms that underlie several forms of strabismus, termed congenital cranial dysinnervation disorders (CCDDs) (Graeber et al., 2013; Whitman and Engle, 2017). Several human mutations that predispose to strabismus have been identified. Testing the function of these genes has been examined in mouse and zebrafish models. These animal models suggest that the waiting zone where oculomotor axons stall, reorient toward muscle targets and initiate branching is a critical period that may be particularly sensitive to disruption (Cheng et al., 2014; Clark et al., 2013; Ferrario et al., 2012; Miyake et al., 2008). Investigation into the role these genes play in guiding the motor nerves has brought renewed attention to not only development of the oculomotor system but mechanisms of neuromuscular connectivity (Chilton and Guthrie, 2017).

Our data demonstrates that the ‘waiting zone’ is a region where the nerve is associated with muscle precursors that are actively signaling to ocular nerves. Therefore, CCDDs that are thought to originate in errors in the ‘waiting zone’ or ‘distal decision zone’ may include defects in guidance cues derived from muscle precursor cells.

## Conclusions

Our study investigates the role that muscle precursors play in motor nerve development in mammals. We find that an important intermediate target of the oculomotor nerve is the mass of eye muscle precursors. Early interaction with muscle precursor cells provides a pause signal to inhibit further growth, and provides a substrate to hold the nerve in a plexus during the previously described ‘waiting period’ or ‘decision region’. Primary myogenesis initiates after muscle precursors migrate away from the mass to positions around the eye. Nerve growth resumes with branching occurring shortly after the formation of myofibers. Ablating muscle precursors early in development results in overshooting the target, while disrupting precursors after plexus formation results in local but aberrant branching of the nerve. This study suggests that muscle precursor cells play an active role in motor nerve development.

## Acknowledgements

Several individuals contributed technical assistance in genotyping, and assistance in optimizing immunohistochemistry including Minkyung Kim, Thomas Kidd, and Hillary Price.

**Author contributions**
BB, GSM, KGW, and TWG formulated ideas for these experiments. Methodology: BB performed nerve tracing, antibody labeling, and analysis; several antibody labels were performed by GER, MV and KGW. LC and PJG contributed Pitx2 mutant embryos and advice on Pitx2 antibody labeling. BB and GSM drafted the manuscript, with editing by KGW and TWG. All authors reviewed and approved the manuscript.

